# Pareto-optimal trade-off for phenotypic switching of populations in a stochastic environment

**DOI:** 10.1101/2022.01.18.476793

**Authors:** L. Dinis, J. Unterberger, D. Lacoste

**Affiliations:** GISC - Grupo Interdisciplinar de Sistemas Complejos and Dpto. de Estructura de la Materia, Física Térmica y Electrónica, Universidad Complutense de Madrid, 28040 Spain; Institut Elie Cartan, UMR CNRS 7502, Université de Lorraine, BP 239 F-54506 Vandoeuvre-lès-Nancy Cedex, France; Gulliver Laboratory, UMR CNRS 7083, PSL Research University, ESPCI, 10 rue Vauquelin, F-75231 Paris Cedex 05, France

## Abstract

Finding optimal survival strategies of living systems embedded in fluctuating environments generally involves a balance between phenotypic diversification and sensing. If we neglect sensing mechanisms, it is known that slow, resp. fast, environmental transitions favor a regime of heterogeneous, resp. homogeneous, phenotypic response.

We focus here on the simplest non-trivial case, i.e. two randomly switching phenotypes subjected to two stochastically switching environments. The optimal asymptotic (long term) growth rate of this model was studied elsewhere; we further expand these results by discussing finite time growth rate fluctuations. An exact asymptotic expression for the variance, alongside with approximations valid in different regimes, are tested numerically in details. Our simulations of the dynamics suggest a close connection between this variance and the extinction probability, understood as risk for the population. Motivated by an earlier trade-off analysis between average capital growth rate and risk in Kelly’s gambling model, we study the trade-off between the average growth rate and the variance in the present model. Despite considerable differences between the two models, we find similar optimal trade-off curves (Pareto fronts), suggesting that our conclusions are robust, and broadly applicable in various fields ranging from biology/ecology to economics.

## 1. Introduction

In unpredictably varying environments, it is advantageous for a population to accept a reduction of its short-term reproductive success in exchange for longer-term risk reduction. This phenomenon, called bet-hedging, protects individuals from potential damages associated with environment variations [1, 2]. It is an important topic in biology which is associated to a number of phenomena such as species polymorphism, antibiotics resistance of bacteria [3] or the resistance of cancer cells to anti-cancer drugs, and more generally to the phenomenon of cell variability [4] and adaptation by the immune system. In all these examples, a dynamic phenotypic heterogeneity at the single cell level brings a fitness advantage at the population level when the environment is fluctuating [5]. Bet-hedging is also a widely studied phenomenon in ecology. For instance, plants use it to delay germination as a form of insurance policy against potentially damaging environment fluctuations [6]. It is important both in spatially homogeneous or heterogeneous environments. In the latter case, it may correspond to a strategy for a given population to colonize an heterogeneous environment [7].

In the literature, an important distinction is made between stochastic bet-hedging, in which the biological system switches stochastically between two phenotypic states at constant rates independent of the environment, and the case of sensing, where the biological system adapts the switching rates to the environment, using information extracted from the environment and relying on a form of memory [8]. The case of adaptive strategies using memory in temporally correlated environments is challenging to describe theoretically but there is constant progress even in this difficult case [9, 10, 11]. In this context, fluctuation relations have been derived for biological populations, which can sense and extract information dynamically from fluctuating environments [12, 13]. These works identified a thermodynamic structure in population dynamics and put forward a deep connection between fitness and information, which underlies the universal adaptation properties of living systems.

Stochastic bet-hedging is perhaps best illustrated theoretically using Kelly’s model, originally introduced in the context of gambling models such as horse races [14]. Kelly proposed a criterion to determine how to place optimally the bets of the gambler so as to maximize the long term growth rate of its capital. The criterion has been used for gambling and for applications in money investment [15]. Being based on information theory, the criterion is general and is also broadly applicable to resource allocation problems in biology, such as the problem of spatial allocation of enzymes within a cell [16]. In practice, Kelly’s strategy is known to be risky, because it implies wild fluctuations of the growth rate of the capital, which most gamblers are not comfortable with. The reason is that Kelly’s model focuses on long term growth but neglects short term risk, which could be very relevant for gamblers and biological populations [17]. A more acceptable solution is an optimization of the mean fitness/growth rate combined with a minimization of the variance, i.e. the risk. In a recent work also inspired by Stochastic Thermodynamics, we have revisited the trade-off between mean growth rate and variance for Kelly’s horse race model, and we have studied the Pareto front formed by the corresponding optimal strategies [18].

In this paper, we go significantly beyond Kelly’s model, by studying a model of a biological population in a fluctuating environment. We assume that the fitness of individuals depends on the environment, and that individuals can switch stochastically between two phenotypic states, at constant rates independent of the environment [19, 20], so that there is no sensing and no memory. We explain why despite these simplifying assumptions, this problem is still considerably more difficult to tackle than Kelly’s original model. To make progress, we introduce a new measure of risk for the population, namely the variance of the finite time growth rate. We first derive an approximate expression of the variance of the finite time growth rate in the limit in which environment fluctuations are slow with respect to phenotypic transitions [8]. Then, we study the general case of arbitrary environment fluctuations and an arbitrary number of discrete phenotypic states, thanks to results derived by one of us in a companion paper [21]. We test both expressions of the variance using numerical simulations in the particular case of two phenotypic states and two environments.

In the literature, many different trade-offs have been considered in this context of populations growing in varying environments. In a classic representation, the growth rate is optimized in the space spanned by the different achievable fitnesses for each separate environment [22, 9]. Another possibility is to look at the distribution of phenotypes in the optimal strategy [23]. Here, we study instead the Pareto-optimal trade-off in terms of the average and the variance of the growth rate. This trade-off is essentially the one between growth rate and risk, which is well documented in economics or in gambling models [15], and which is also relevant for biological and evolutionary processes [24, 25, 26, 27]. Using numerical simulations, we also show that the variance of the growth rate is an acceptable measure of risk for the population, because strategies with a high growth rate variance are the ones with a higher probability of extinction.

## 2. Definition of the model

Let us consider a biological population of individuals which exhibit only two phenotypes *A* and *B*, which can randomly switch between them. To simplify let us also assume that the environment has only two discrete states 1 and 2 [5, 2]. We denote the population vector, which describes the number of individuals in each phenotype (*A* or *B*) at a given time *t* by **N**(*t*) = (*N_A_*(*t*), *N_B_* (*t*))^*T*^, where *T* denotes the transpose. The subpopulation of individuals with phenotype A grows when placed in the environment *i* with the growth rate *k_Ai_*, while the other subpopulation with phenotype B grows with rate k_Bi_. There is no population noise, the dynamics of the system is deterministic in each separate environment and individual growth rates can take positive or negative values [8].

When both growth rates take positive values, the evolution of the two subpopulations is equivalent to that of two species (also called A and B), which grow according to autocatalytic reactions. The corresponding chemical reactions are

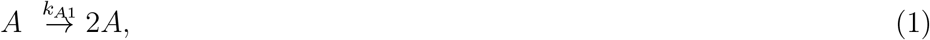

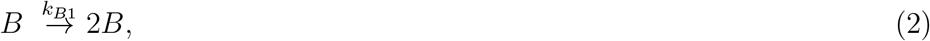

for the growth of the phenotypes (A, B) in environment 1 and similarly,

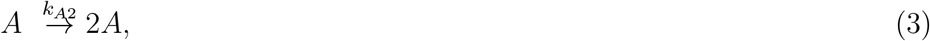

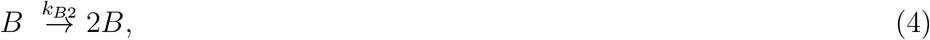

for the growth in environment 2. Environmental transitions, which are stochastic, can be described by the reversible reaction:

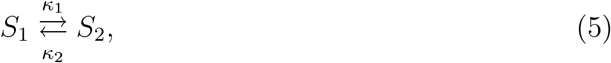

where *S*_1_ (resp. *S*_2_) represents environment 1 (resp. 2).

For applications, we shall assume in addition that phenotype *A* is more adapted to environment 1 than phenotype *B*, so that *k*_*A*1_ ≥ *k*_*B*1_; while phenotype *B* is more adapted to environment 2, so that *k*_*A*2_ ≤ *k*_*B*2_ [19]. Let 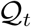 be the marginal probability of the environment at time *t.* Since the evolution of the system and environment states form a Markov process in continuous time, this probability distribution admits the stationary measure defined by 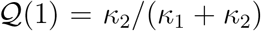, for the probability of the environment to be in the first state and 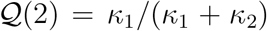 for the other state. The different periods of environment *i*, denoted *τ_i_* are assumed to be i.i.d. exponentially distributed random variables.

Independently of the state of the environment, individuals can switch their phenotype. These phenotypic transitions can be described chemically by the reversible reaction:

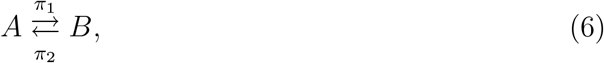

which is always present irrespective of the state of the environment. These rates *π*_1_ and *π*_2_ represent the strategy of the individual, similar to the betting strategy in Kelly’s horse races. Note that there is no sensing, which means that these rates are independent of the state of the environment.

All these reactions can be summarized by the vector equation

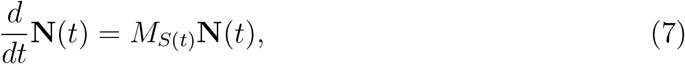

with matrices

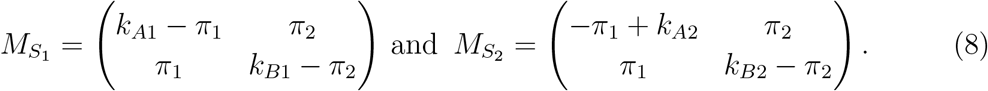

The finite time averaged population growth rate is defined as

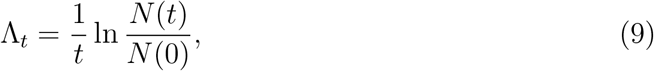

in terms of the total population *N*(*t*) = *N_A_*(*t*) + *N_B_*(*t*), and the long term population growth rate is

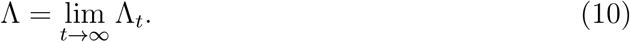

When the environment remains constant, i.e. when *S*(*t*) = *S_i_* for all times, the total population grows exponentially with a growth exponent equal to the top eigenvalue of the matrix *M_S_i__*, while the distribution of phenotypes is determined by the corresponding eigenvector, denoted **q**_*i*_.

### 2.1. Main quantities of interest

In the general case of a switching environment, it is more difficult to obtain an analytical expression of the growth rate, because one needs to evaluate a product of a large number of random matrices of the type

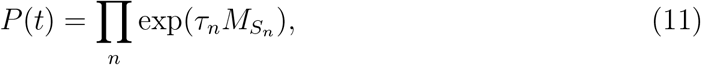

where the product is over the various alternating environments of duration *τ_n_* such that *Σ_n_ τ_n_ = t.* The quantity we are interested in is called the Lyapunov exponent in the literature, which corresponds precisely to the growth rate defined previously:

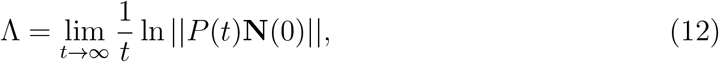

an expression which is known to be independent of the choice of norm denoted || · || for the matrices and independent of **N**(0), an arbitrary vector describing the initial condition [28]. Another important property of that Lyapunov exponent is that it is a self-averaging quantity, therefore there is no need to average over the ensemble of random matrices: Λ = 〈Λ〉. Although there is no simple method to compute that Lyapunov exponent exactly in the general case where the matrices do not commute (except in the case of 2 x 2 matrices as done in [19]), there are a number of useful approximations, which generalize to arbitrary dimensions.

For real application, demographic fluctuations are important because in the end, one is always interested in finite populations in a finite time [17, 24]. These effects cannot be predicted from A alone; one should consider instead the finite time growth rate Λ_*t*_ and its fluctuations characterized by the variance Var(Λ_*t*_). To evaluate this variance numerically, one needs to carry out a sufficiently large number of independent simulations, all starting with the same initial conditions. A quantity similar to the variance Var(Λ_*t*_) (and higher moments too) has been considered in the mathematical literature on large products of random matrices [28, 29].

Another important quantity in this context is the instantaneous growth rate *μ*(*s*), defined as

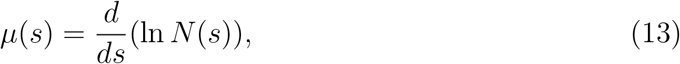

so that Λ_*t*_ reads:

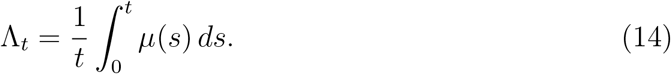

Since instantaneous growth rates decorrelate exponentially fast, the central limit theorem imposes a scaling of Var(Λ_*t*_) in 1/*t* as *t* → ∞. Therefore, our main focus is the evaluation of lim_*t*→∞_ *t*Var(Λ_*t*_), a self-averaging quantity, which we denote (by abuse of notation)

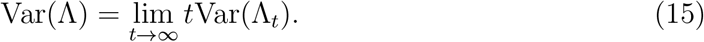

## 3. Kussel-Leibler approximation

In the so-called adiabatic approximation, one assumes that environment periods are long enough so that the population has time to reach an equilibrium distribution (given by the top eigenvector in that environment) before the environment switches again. Such an approximation was introduced by Kussel-Leibler (KL) to evaluate the long term population growth rate in a fluctuating environment and the optimal phenotypic strategy, in terms of the characteristic switching dynamics of the environment [8].

### 3.1. Mean growth rate

Their general expression of this long-term growth rate in the particular case of two environment states and two phenotypic states takes the form:

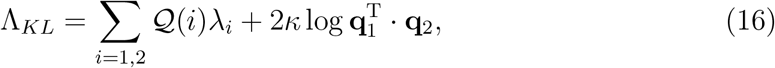

where λ_*i*_ is the top eigenvalue of the matrix *M_s_i__*, **q**_*i*_ the corresponding top eigenvector; *κ_i_* = 1/〈*τ_i_*〉, *i* = 1, 2 are the inverse of the average periods of each environment; and 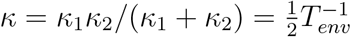, where 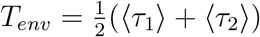 is the average time span of an environment.

The first term in the r.h.s. of Eq. 16 corresponds to the average growth rate, where the average is taken with respect to the stationary measure 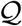, which is equivalent to an average over the fractions of times spent in each of the two states, in the limit where these times become infinite. The second term in the r.h.s. of Eq. 16, which is negative, is a penalty due to transitions between the two environments. This term features the overlap between the two dominant eigenvectors, which arises due to the change of base in going from the top eigenvector of one environment to the top eigenvector of the other. For this reason, this term depends on 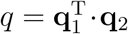; it would vanish if the two matrices *M*_*S*_1__ and **M*_*S*_2__* commuted. In practice however, this is never the case, and this contribution due to the change of basis is the main reason for the difficulty in obtaining an exact expression of the grow rate.

Simple explicit formulas follow from a Taylor expansion in the case where the switching rates *π_i_* are small compared to the differential growth rates |*k_A_i__* — *k_B_i__*|. Assuming that the growth matrix is diagonal, i.e. that only one phenotype grows in one environment but not in the other, in other words when *k*_*A*1_ = *k*_1_ > 0, *k*_*B*1_ = 0, *k*_*A*2_ = 0 and *k*_*B*2_ = *k*_2_ > 0, the top eigenvalues for the two environments *i* =1, 2 are *λ_i_* ≃ *k_i_* – *π_i_* to first order in *π_i_/k_i_*, and *q* ≃ *π*_1_*π*_2_/*k*^2^, where *k* = *k*_1_*k*_2_/(*k*_1_ + *k*_2_). Therefore, when *k_i_* » *π_i_*, the above expression simplifies into:

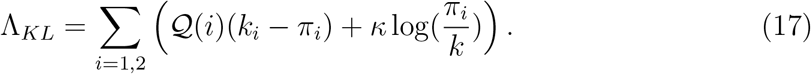

In addition to the condition *k_i_* ≫ *π_i_*, the KL approximation requires that the second term in Eq. 16 be small with respect to the first term, which leads to the condition log(*k/π_i_*) ≪ *k/κ*. In the case where *k*_*B*1_ and *k*_*A*2_ are not zero, this criterion is still approximately correct provided one uses for *k*_1_, resp. *k*_2_, the relative growth rate *k*_*A*1_ – *k*_*B*1_, resp. *k*_*A*2_ — *k*_*B*2_.

By optimizing Λ_*KL*_ with respect to *π_i_*, one finds

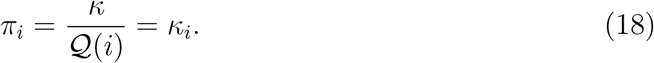

Thus, the optimal strategy corresponds to switching rates that match the environment rates. By reporting these optimal transition rates in Eq. 17, one finds that the optimal growth rate is

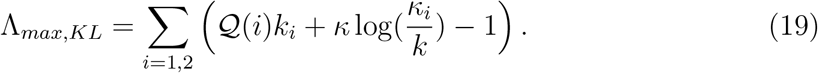

It is easy to see that this growth rate is maximum [8], because

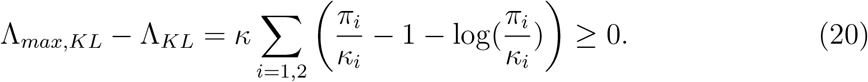

This condition of optimality *π_i_* = *κ_i_* is very similar to Kelly’s criterion [14], which leads to the maximum of the capital growth rate in the betting game. As shown by Kussel-Leibler, this condition remains true in the general case where all the growth rates take non-zero values.

### 3.2. Variance of the growth rate

Within the KL approximation, the growth rate in the limit of a large number of environmental epochs only depends on the fraction of time spent in the first environment *r* and on the total number of transitions 2*N* between the two states. The joint distribution of *r* and *N*, namely *f* (*r,N*) is easily expressed in terms of the product of two Poisson distributions of parameters *κ*_1_*rt* and *κ*_2_(1 – *r*)*t*. One can check that this distribution is maximum when *N* ≃ *κt* and *r* ~ *Q*(1). Then, we rely on a Gaussian approximation of that distribution close to the maximum to evaluate the variance, which becomes more and more accurate as *N* becomes large. Details of this calculation are provided in appendix A. We find that for large *t*, under the same approximations leading to (17):

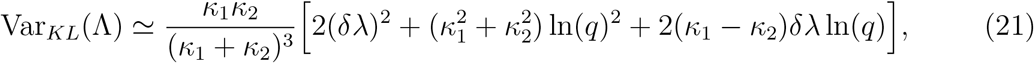

where *δλ* = λ_1_ – λ_2_.

In the section on numerical results 6.1, this expression will be tested and compared with other expressions of the variance of the growth rate.

## 4. Approximation for fast environmental changes

We now study a different approximation, which is in some sense opposite to the adiabatic approximation considered in the previous section, namely the approximation of fast environmental changes. Below, we use the piecewise-deterministic Markov process (PDMP) rewriting of the model introduced by Hufton et al. [20], and focus on the case of two environment states and two phenotypic states [19] where explicit computations are available. For a more general discussion of the PDMP at high frequency, where discrete transitions are fast, we refer the reader to Ref. [30].

### 4.1. Average growth rate

Hufton and Lin introduced the relative fraction of phenotype *A* in the population *ϕ* = *N_A_*/(*N_A_* + *N_B_*), which evolves deterministically in each environmental epoch according to a differential equation. Fixed points of the dynamics are denoted 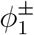 in the first environment and similarly 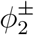 in the second one; explicitly,

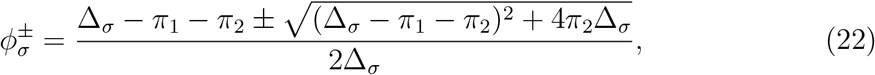

with Δ_*σ*_ = *k_Aσ_* — *k_Bσ_* for *σ* = 1,2. The + superscript indicates the solution which is a stable fixed point, while the – superscript denotes the unstable one. As discussed in the introduction, we assume that phenotype *A* is more adapted to environment 1 than phenotype *B*, so that *k*_*A*1_ ≥ *k*_*B*2_; while phenotype *B* is more adapted to environment 2, so that *k*_*A*2_ ≤ *k*_*B*2_; this means that Δ_1_ ≥ 0 and Δ_2_ ≤ 0 [19].

The stationary probability density distribution *P_σ_* (*ϕ*) of the Markov process describing the evolution of the relative fraction *ϕ* in a stochastically switching environment can be solved exactly using the method of characteristics. The solution has support on 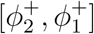; it depends on the two switching rates *κ_σ_* and on the fixed points 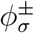. With these notations, the stationary probability distributions read:

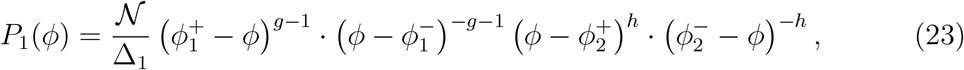

and

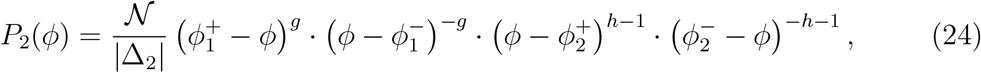

where *g* and *h* are positive exponents given by

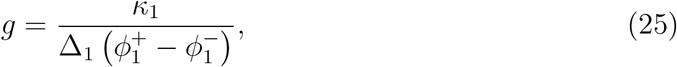

and

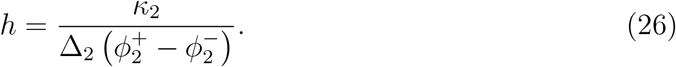

The integration constant 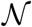 is fixed by the normalization condition

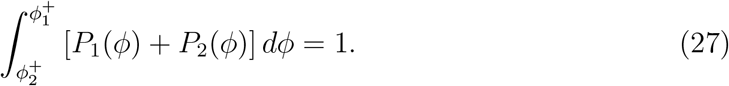

Once that value of 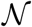 has been determined, one obtains the two separate relations

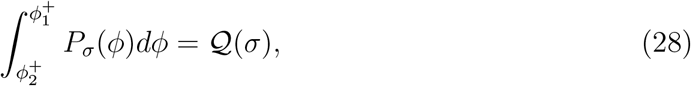

for *σ* = 1, 2, which may be view as the marginal distribution in *σ* of the joint distribution *P_σ_* (*ϕ*) over *ϕ.*

Note that *P*_1_ contains a singularity at 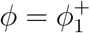, and similarly for *P*_2_ at 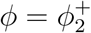. As a result, integrals involving *P*_1_ and *P*_2_ can be difficult to evaluate numerically. Fortunately, that difficulty can be overcome by using a change of variable, 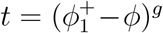 with integrals involving *P*_1_ and 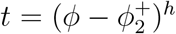 for those involving *P*_2_. For instance, the integral of Eq 28 for *σ* = 1 is turned with this change of variable into the following integral free of singularity:

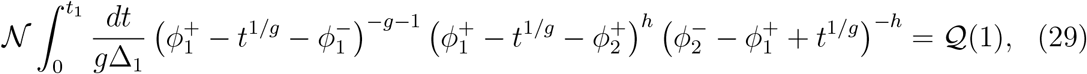

where 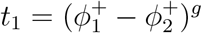.

This trick is useful to evaluate the normalization constant 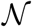 but also the following two integrals *K*_1_ and *K*_2_, from which the average growth rate Λ can be obtained. The two integrals are

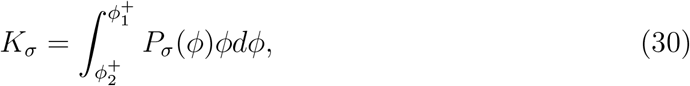

for *σ* = 1, 2.

Hufton and Lin also introduced the instantaneous growth rate, which they define as [19]:

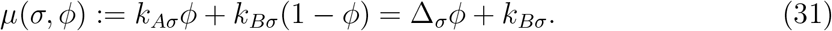

and mention that it is possible to obtain an exact expression of the stationary distribution of *μ* = *μ*(*σ,ϕ*), although they do not give it explicitly. Here is how we obtain it. Let 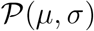 be the stationary distribution of the instantaneous growth rate *μ* in the environment *σ*. This distribution can be obtained from that of *P_σ_* by changing variables from *ϕ* to *μ* at fixed value of the environment *σ*. One obtains 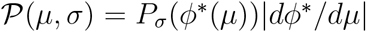, where *ϕ*\*(*μ*) = (*μ* – *k_B_σ__*)/Δ_*σ*_ is the function that inverts Eq. 31. Thus, we obtain

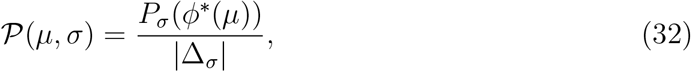

with support 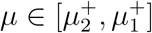, where 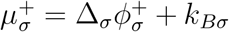. The distribution of *μ* can then be obtained by marginalizing over *σ*:

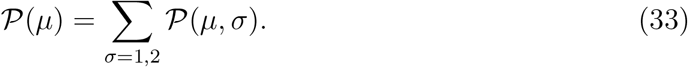

This distribution is smooth when the environment changes quickly, but contains singularities in the general case as shown in Fig 3 of [19].

The average growth rate of Eq. 35 is obtained from the first moment of that distribution Λ = 〈*μ*〉, because of the following equalities:

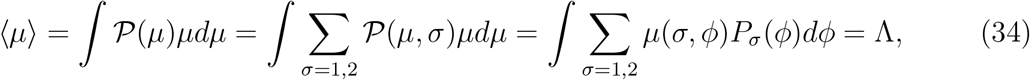

where *μ*(*σ, ϕ*) is the function defined in Eq. 31. The average growth rate can then be written explicitly in terms of the integrals introduced above as

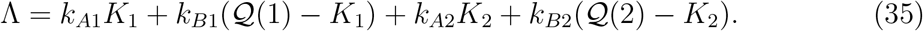

### 4.2. Variance of the instantaneous growth rate

The second moment of that distribution 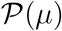 represents the variance of the instantaneous growth rate, which can be easily obtained in this framework:

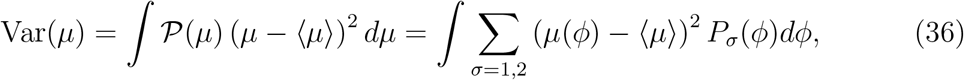

Explicitly, we have

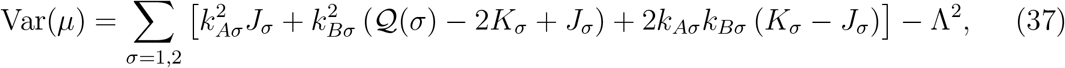

which depends on additional integrals of the form

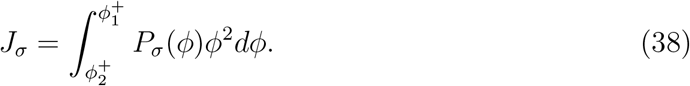

All these integrals can be written in a closed form, free of divergences, by using the same trick introduced above.

## 5. Exact solution

It is important at this point to appreciate the difference between the finite time growth rate variance Var(Λ_*t*_) and Var(*μ*). Recall that 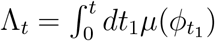, where *ϕ*_*t*_1__ is the fraction of phenotype *A* in the population at time *t*_1_. It then follows that

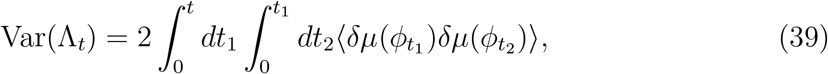

with *δμ* = *μ* — 〈*μ*〉 and the function *μ*(*ϕ*) is defined in 31. This expression makes clear that time correlations of the instantaneous growth rate contribute to Var(Λ_*t*_), but not to Var(*μ*). This is also the reason why in practice the instantaneous growth rate distribution 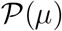 is found to be in general different from the distribution of Λ_*t*_ [19].

To address this issue, we use an exact expression of the asymptotic behavior of Var(Λ_*t*_) derived by one of us by a PDE approach in a companion paper [21], which contains the approximations introduced above as particular cases. We mention here only the final and main result of this work. In the framework introduced by Huffton and Lin, this solution takes the following form: let

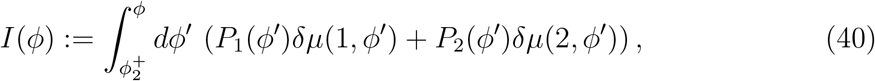

and

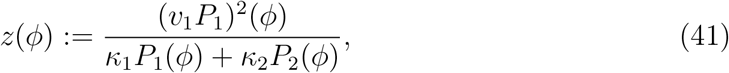

where *v*_1_(*ϕ*) = Δ_1_*ϕ*(1 – *ϕ*) – *π*_1_*ϕ* + *π*_2_(1 – *ϕ*).

Then, the asymptotic behavior of the finite time growth-rate variance *t*Var(Λ_*t*_) at large times *t* is

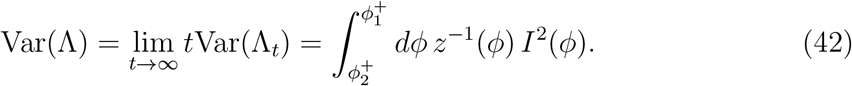

This expression is tested in the next section with numerical simulations.

## 6. Numerical results

### 6.1. Growth rate as a function of the rate of change of the environment

In fig. 1, we show as a heatmap, the average growth rate for different switching rates of the environment. We used the set of parameter values *k*_*A*1_ = 2, *k*_*B*1_ = 0.2, *k*_*A*2_ = – 2, *k*_*B*2_ = —0.2, which correspond to the ones used by Hufton and Lin in their figure 4 [19].

**Figure 1:**
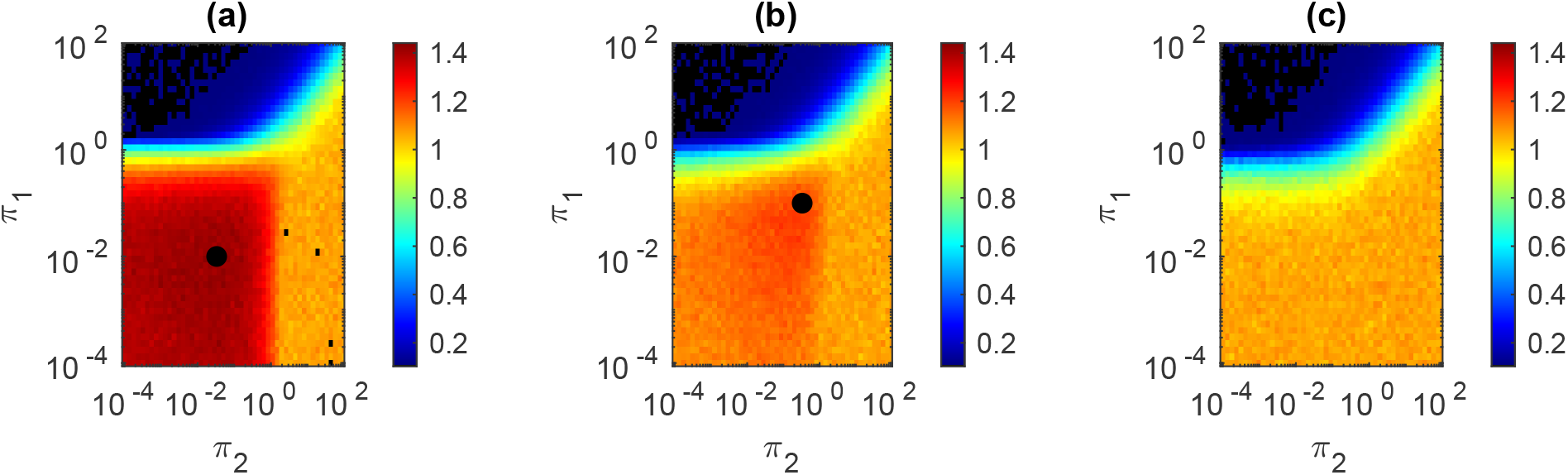
Heatmap plot of the average long term growth rates for different values of the phenotypic switching rates *π*_1_ and *π*_2_. Figure (a) corresponds to *κ*_1_ = 0.01, *κ*_2_ = 0.03, (b) to *κ*_1_ = 0.1, *κ*_2_ = 0.3, (c) to *κ*_1_ = 1, *κ*_2_ = 3.3. The black dot in Fig (a) and (b) indicate the point where the average growth rate takes its maximum value according to the KL approximation.

Three different switching rates have been used for environmental fluctuations, for case (a): *κ*_1_ = 0.01, *κ*_2_ = 0.03, for case (b): *κ*_1_ = 0.1, *κ*_2_ = 0.3, for case (c): *κ*_1_ = 1, *κ*_2_ = 3.3. In each case, about 50 simulations of duration 100/ *κ*_1_ have been performed. In the case of Fig 1a and b, for a slow switching rate, the maximum value of the growth rate is reached on isolated points in this diagram, i.e. for specific values of *π*_1_ and *π*_2_. These values correspond well to the condition *π_i_* = *κ_i_* predicted by the KL approximation, where the phenotypic switching rates match that of the environment, which is represented by a black dot on the figure.

Since the fastest growth rate is *k*_*A*1_, one could expect that the highest growth rate should be obtained when the system spends most of the time with the phenotype *A*, and the lowest growth rate when it stays with the phenotype *B.* The latter hypothesis is confirmed by Fig. 1a, because the smallest value of the growth rate is indeed obtained in the top left part of the figure, i.e. when *π*_1_ is large and *π*_2_ is small, which corresponds to conditions where the subpopulation with phenotype *A* turns instantaneously into the phenotype *B*. The former hypothesis however is not confirmed, because the fastest growth rate is not obtained for large *π*_2_ and finite *π*_1_, in that particular case, it is obtained when both *π*_1_ and *π*_2_ are small, i.e. in the KL regime.

When the environment switches very fast, there is no isolated maximum in these heatmap plots as shown in Fig 1c, in that case the optimum growth rate is reached on the boundaries of the simplex in which *π*_1_ and *π*_2_ take their values. Thus, phenotypic heterogeneity presents a fitness advantage only for slow environments (cases a and b), which are accessible to the KL approximation. In contrast, when the variations of the environment are fast (case c), phenotypic homogeneity is favored, which is a regime beyond the validity of the KL approximation [19].

### 6.2. Validity of the various approximations

To check the various approximations more precisely, we compare in table 1 the average and the variance of the growth rate obtained from simulations with their estimations based on various approximation schemes. We used the same values of *k*_*A*1_, *k*_*B*1_, *k*_*A*2_, *k*_*B*2_ as above, together with four new sets of environmental and phenotypic transition rates, which we call (d), (e), (f) and (g). The parameters are, for case (d): *κ*_1_ = *κ*_2_ = 0.1 and *π*_1_ = *π*_2_ = 0.064, case (e): *κ*_1_ = *κ*_2_ = 1 and *π*_1_ = *π*_2_ = 0.24 and case (f): *κ*_1_ = *κ*_2_ = 10 and *π*_1_ = *π*_2_ = 0.4, case (g): *κ*_1_ = *κ*_2_ = 0.01 and *π*_1_ = *π*_2_ = 0.064.

**Table 1:** Comparison between different estimations of the average and variance of the growth rate for four data sets. The star indicates results which are not meaningful because the assumptions needed for the approximation are not met. In this table, Λ_*KL*_ corresponds to the theoretical growth rate evaluated from Eq. 16, Var(Λ) has been evaluated from Eq. 42, Λ is obtained from Eq. 35, Var(μ) from Eq. 37 and finally Var_*KL*_(Λ) from Eq. 21

| Data | Case d | Case e | Case f | Case g |
| --- | --- | --- | --- | --- |
| $\Lambda$ | 0.638 | 0.238 | 0.037 | 0.81 |
| $\text{Var}(\Lambda)$ | 12.5 | 1.3 | 1.28 | 120.4 |
| $\text{Var}(\mu)$ | 1.4 | 1.37 | 1.24 | 1.24 |
| $\Lambda_{KL}$ | 0.57 | * | * | 0.81 |
| $\text{Var}_{KL}(\Lambda)$ | 12.27 | * | * | 121.07 |

The average growth rate Λ has been measured using numerical simulations, which have been found to agree with Eq. 35. This confirms that correlations of the instantaneous growth rate do not matter for the average growth rate. The KL approximation is found to provide a good estimate of the average and variance of the growth rate when *π*_1_, *π*_2_, and *κ* are small compared to the growth rates of the phenotypes in their respective environments, conditions which are satisfied for case (d) and (g) only. In contrast for cases (e) and (f), the KL approximation breaks down and fails to provide estimates for the average and variance of the growth rate.

In the regime of fast environment changes for cases (e) and (f), the variance of the finite time growth rate agrees well with the variance of the instantaneous growth rate, which is to be expected since the environment time correlations are very short compared to other time scales.

For the general case, we have also checked that the theoretical expression of the variance of Eq. 42 gives correct results in all cases (d) to (g). To illustrate this point further, we provide an additional figure Fig 2 corresponding to the specific parameters of case (e). In that figure, we compare the analytical expression with numerical simulations for various duration times t. In practice the average of the variance is evaluated from a number of independent simulations, whose number is also proportional to t. As shown in the figure, there is a very good agreement provided the time t is sufficiently long. The duration of that initial transient depends on the number of simulations as expected.

**Figure 2:**
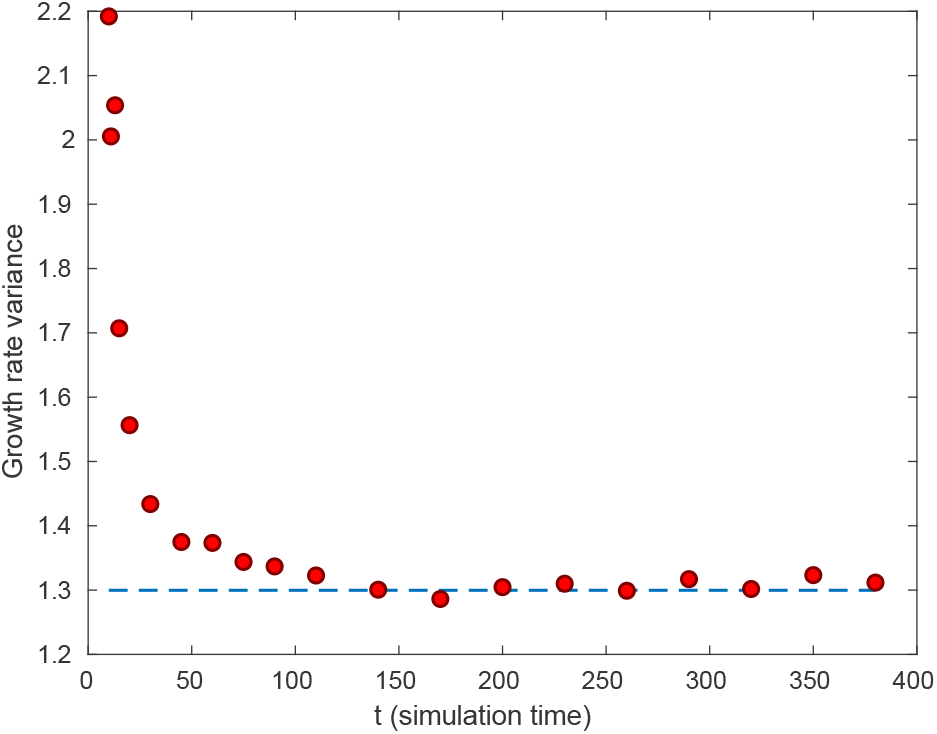
Growth rate variance as function of simulation time *t* for parameter set (e), with symbols corresponding to the numerical simulation and the dotted line corresponding to the asymptotic theoretical prediction of Eq. 42.

### 6.3. Pareto fronts

We now analyze the relation between mean growth rate Λ and the asymptotic behavior of the finite time growth rate variance, which we denoted Var(Λ). As stated before, higher growth rate can lead to higher fluctuations (or risk) and therefore a suitable balance between average growth rate and variance may be advantageous. As in previous sections, the values of (*π*_1_,*π*_2_) constitute the strategy of the individuals (or colonies) for given environmental parameters. The optimal trade-off is given by the maximum growth attainable for a fixed level of fluctuations, or conversely, by the minimum variance possible for a fixed mean growth rate. The 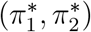 that optimize the trade-off can be found by minimizing the following objective function, which is a a linear combination of both quantities

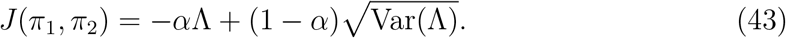

In this objective function, the standard deviation is used as a measure of risk instead of the variance in order to keep the risk tolerance parameter *α* dimensionless. Note that the Pareto front and the trade-off are not affected by this choice, which means that these features should be similar across different systems.

Minimization of function *J* for different *α* has been performed by a simulated annealing algorithm. Starting from initial values for (*π*_1_,*π*_2_) a random move in this 2D space is either accepted if it decreases the objective function J, or accepted with an exponentially decaying probability if it increases J. The exponential probability is controlled by an effective temperature parameter that is progressively decreased (hence annealing) making it harder and harder to accept an increasing move. These upward moves allow the algorithm to escape local minima initially and proceed to the global minimum.

Once the optimal values 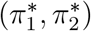 are obtained, one can compute the values of Λ and 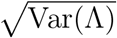 to which they correspond, thus building the efficient border or Pareto front, represented in figure 3a. Any strategy on that front cannot be improved in terms of one property (average or variance) while keeping the other constant, and therefore represents the optimal trade-off. Some of these strategies are represented as colored dots in the figure.

**Figure 3:**
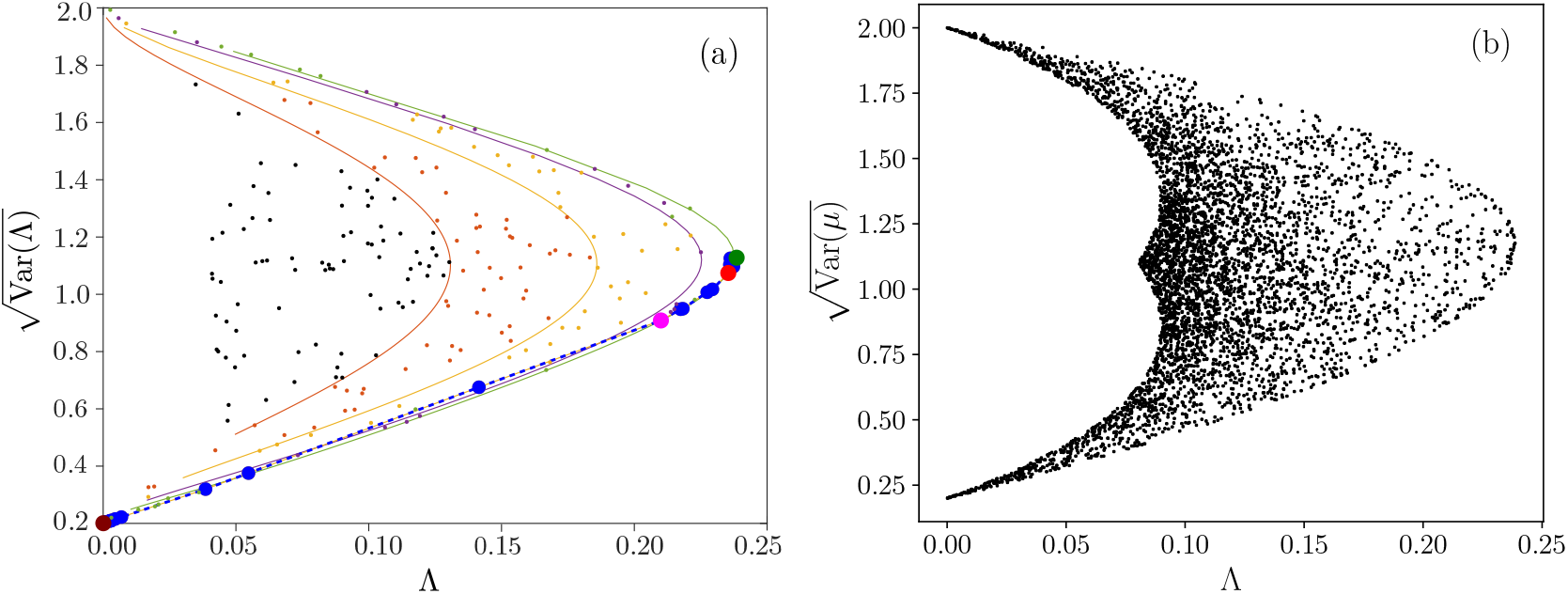
Mean-variance trade-off for the long term growth rate Λ (in (a)) or for the instantaneous growth rate (in (b)) using parameters as in set (e) except for (*π*_1_,*π*_2_) which are varied. In figure (a), filled (blue or other color) circles represent points in the Pareto front computed by minimizing the objective function *J*(*π*_1_,*π*_2_; *α*). The dashed blue line interpolates the front between the computed points. For four highlighted points in the Pareto front, marked with green, red, magenta and maroon filled circles, we provide their coordinates in table 2. Solid thin lines: constant T lines, from left to right *T* = 0.5 (orange), *T* =1 (yellow), *T* = 2 (violet) and optimal *T* = 3.33 (green). Colored dots are obtained by scanning (*π*_1_,*π*_2_), and the colors are chosen according to the corresponding *T* value. Black dots have *T* < 0.5, orange dots have 0.5 < *T* < 1, yellow dots have a 1 < *T* < 2 and so on.

Environmental and phenotypic changes are associated with two characteristic time scales *T_env_* and *T*. The first time scale 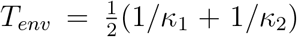 has been introduced in section 3 and represents the average time span of an environment, while the second time scale 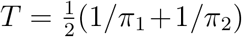 characterizes phenotypic changes. We conjecture that the ratio of these two time scales is a key parameter for the study of the growth rate, and that optimal average growth rates are found when this ratio is close to one. Indeed, the Kussell-Leibler optimum (18) is obtained when the two time scales are of comparable order of magnitude: *T_env_* ~ *T*. This hypothesis is confirmed by plotting curves of constant ratio *T/T_env_* in the plane of the mean growth rate and the standard deviation as shown in Figure 3a. We then observe that all the curves converge to the right as this ratio goes to 3.3, eventually reaching the Pareto front when the ratio approaches 3.3.

In figure 3b, we build a similar diagram for the instantaneous growth rate instead of the long term growth rate. We observe that the right border of the cloud of points, which forms the Pareto front has a similar shape as before. Indeed, with the chosen parameters, 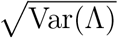 is numerically close to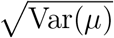, although this is of course not always the case as shown in Table 1.

In both Pareto fronts, fluctuations of the growth rate become small when the average of the growth rate also becomes small, similarly to what happens on the tradeoff branch in Kelly’s model [18]. In that model, the origin of the diagram corresponds to a ‘null strategy’ where both the mean and the variance vanish. Here the origin does not belong to the front, but in fact it does not matter because the absolute value of the mean growth rate is not meaningful, only differences of the growth rate with respect to some reference are significant.

Further, near the point of maximum growth rate, which is similar to Kelly’s point, the slope of the front appears nearly vertical similarly to what we found in our previous study of Kelly’s gambling [18]. This means that, by moving slightly along the front away from this point, fluctuations can be decreased significantly without a large loss of average growth rate loss. We address the significance of this statement by considering the risk of extinction in the following section.

### 6.4. Extinction

As an illustration of the mean-variance trade-off, we now include extinction in our model and check whether larger fluctuations may indeed increase the probability of extinction as implied in previous sections. We will compare Kelly’s point (giving optimal average growth rate, in green in figure 3) against another point in the Pareto front (sub-optimal, in red) giving slightly less growth rate but significantly lower variance, due to the high slope of Pareto front. The actual predicted values computed with the theoretical expressions are given in table 2:

**Table 2:**
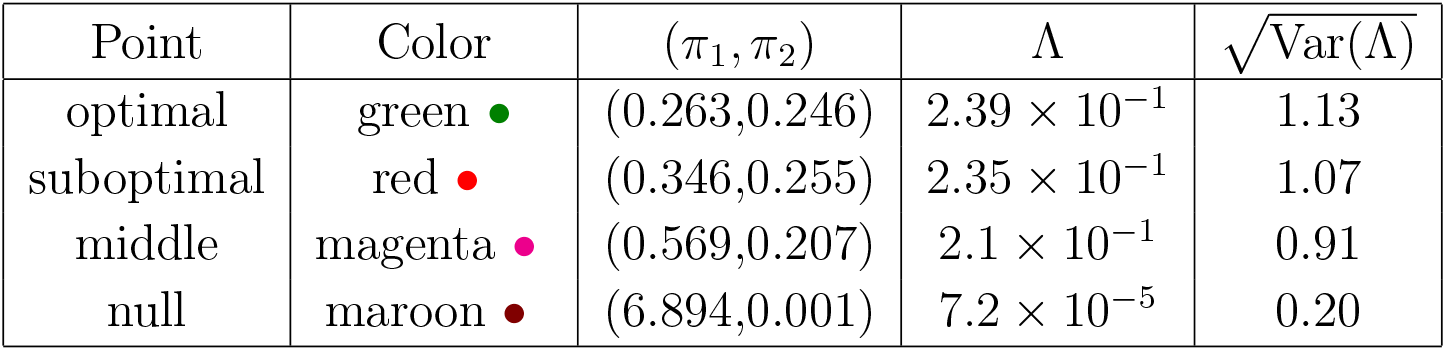
Some points in the Pareto front. Colored circles correspond to the symbols used in figure 3 to indicate specific points.

Points only differ in the corresponding π values, the rest of the parameters are equal *k*_*A*1_ = 2,*k*_*B*2_ = — 0.2, *k*_*A*2_ = — 2, *k*_*B*1_ = 0.2 and *κ*_1_ = *κ*_2_ = 1.0. We run 8000 simulations of evolution equations (7) with each set of parameters for a moderate time *T_max_* = 500/*κ*_1_. The inset in figure 4 depicts the trajectory of log *N*(*t*), in green the ones corresponding to optimal growth rate, the red corresponding to sub-optimal. In the figure one can see that both growth rates are very similar as expected and that fluctuations are slightly more intense in the optimal case. This can be checked by averaging the growth rate in different realizations. Notice that 8000 trajectories were depicted for each case although most of the green trajectories are not visible below the red ones. Nevertheless, several extreme green trajectories are still visible around the borders and cover a wider area indicating higher variance.

**Figure 4:**
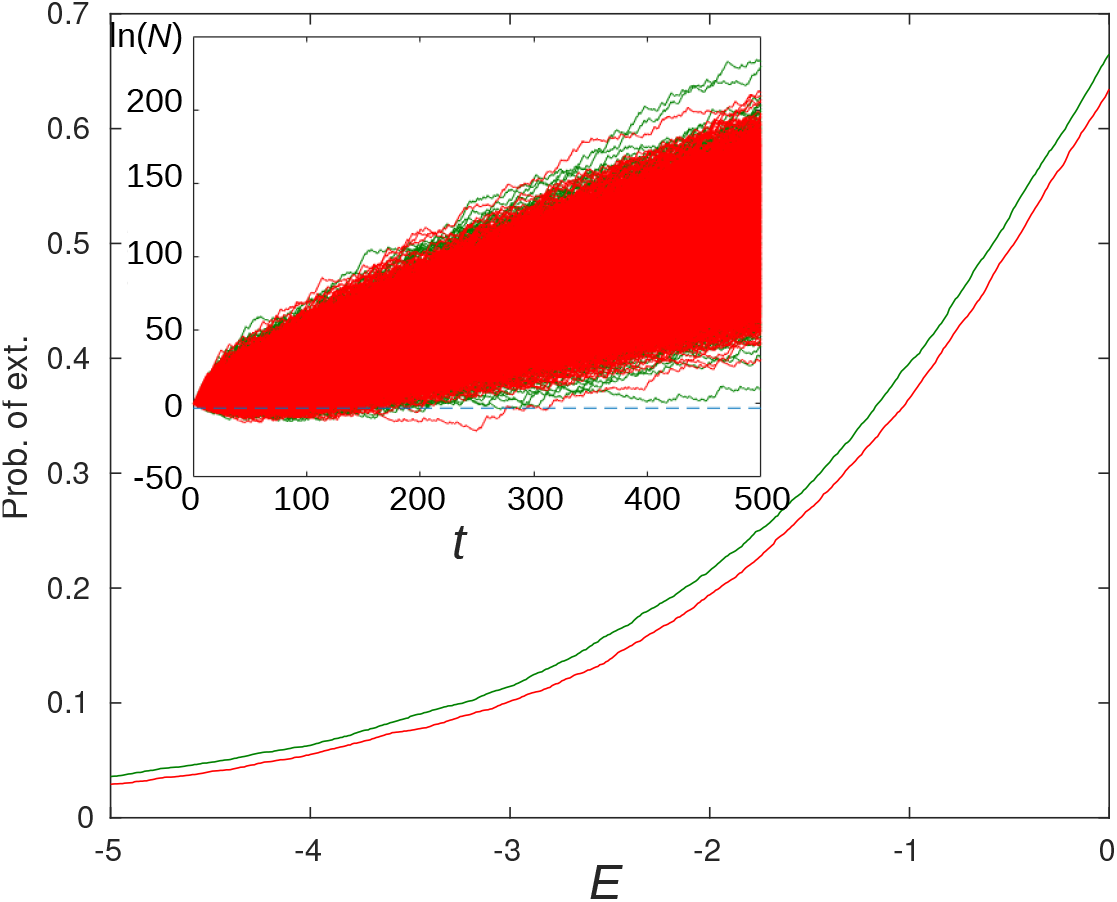
Probability of extinction as a function of threshold *E* for extinction for optimal set of parameters (green) and sub-optimal (red). Inset: simulated trajectories for the optimal set of parameters (green) and sub-optimal (red) and threshold value *E* = —3 (blue dashed line). All trajectories crossing the threshold at any time are considered extinct.

We now set a threshold for extinction *E* < 0 on the logarithm of the population. If the trajectory of log N goes below this threshold at any time t during simulation, the population is considered extinct in this realization. By computing the fraction of realizations that would go below a given threshold, we estimate the probability of extinction for both parameter sets as shown in figure 4. To describe extinction exactly, a stochastic approach would be necessary but for all practical purposes, we consider a population of sufficiently low values of *N* as extinct. We fix the initial population to be equal to one for all realizations.

As expected, the greater the distance between initial population and threshold, the lower the extinction probability. We observe that the probability of extinction is higher for the optimal case than for the sub-optimal one. In the presence of extinction, a colony with smaller growth rate could achieve higher fitness as measured by a lower extinction probability due to its lower variance [31]. In this case, the successful colony trades some growth rate for less risky fluctuations.

However, going further away from the optimum on the lower branch of the Pareto front, the probability of extinction raises again, as checked with the maroon point in figure 3. This confirms the existence of a trade-off between the growth rate and the variability.

## 7. Conclusion

Kelly’s original paper contained two insights, the idea of the optimization of the long term growth rate and its information theoretic interpretation. Despite its remarkable successes in fields ranging from gambling to biology, Kelly’s model is limited because it focuses on the long term growth rate and misses the short term risk, which is relevant to gambling where it can cause ruin of the gambler and to biological populations where it can lead to extinction.

To address this issue, we have studied the variance of the finite time growth rate, which needs to be distinguished from the instantaneous growth rate, because the later is less relevant for the evolution of biological systems. In the case of two environments and two phenotypes, we have derived various approximations for this quantity and tested with simulations an exact, albeit complicated, expression valid for arbitrary durations of the environment fluctuations.

Using this variance, we have built the corresponding Pareto front which characterizes the trade-off between the average growth rate and the risk. We found that this trade-off has similarities with the one we had analyzed previously in our work on Kelly’s model [18], suggesting a form of universality for this trade-off. We have also shown that the risk measured from the variance is indeed linked to the extinction probability of the population. The known experimental observation that bacterial populations faced with stressful conditions maintain a fraction the population with a reduced growth as a form of ‘insurance policy’ to avoid extinction [3] is compatible with this trade-off.

It would be interesting to explore further extensions of our framework to cases where sensing is present and where more phenotypic states are available. As a first step towards including sensing, one of us recently studied adaptive strategies in Kelly’s model [32]. If these ideas can be extended to the problem of populations facing unpredictable environments, one may obtain from them an understanding of the adaptation process at the information level, comparable to what has already been achieved for gambling models. In practice, another complication arises in biological populations, namely that diversification occurs both at the cellular level and at the population level; taking both features into account in the same model will require yet another level of extensions of the present framework.

We hope that in the future, quantitative predictions of our model could be tested experimentally. Experiments on growing colonies with bacteria [33] or with yeasts [34] hold great potential for this kind of tests, because on one hand, cell populations can be monitored continuously on long times, and on the other hand, a fluctuating environment (either periodic or stochastic) can be imposed externally on the system in a controlled way.

## 8. Acknowledgements

We acknowledge many fruitful discussions with O. Rivoire and N. Desprat, and A. Despons for a careful reading of the manuscript. L.D. acknowledges financial support from Spanish Ministerio de Ciencia e Innovación through grant PID2020-113455GB-I00. DL acknowledges support from (ANR-11-LABX-0038, ANR-10-IDEX-0001-02).

## Appendix A. Variance in the KL approximation

### Appendix A.1. Details on the derivation

Here, we provide a derivation of the formula of Eq. 21 for the variance in the KL approximation. In the limit of a large time *t*, the two unknowns in this problem are the number of transitions 2*N*, and the fraction of time spent in the environment state *S*_1_, which we denote *r*. Since 0 < *r* < 1, 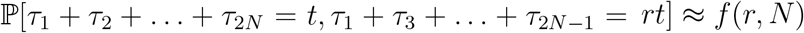, where

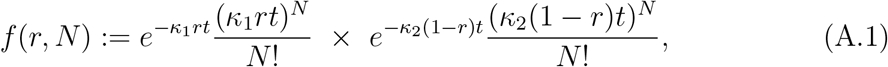

corresponding to the product of two Poisson distributions of parameters *κ*_1_*rt* and *κ*_2_(1 – *r*)*t*. When *t* is large, we have asymptotically in terms of the top eigenvalues *λ_i_* of matrices *M_S_i__*:

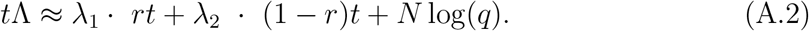

(1) Let us first check that *f* (*r, N*) is maximum when 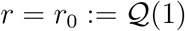 and *N*:= *N*_0_ ~ *κT*. Note that 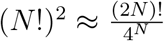, and the function 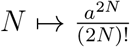 is maximum, equal to ≈ *e*^2*N*^ for *a* ~ 2*N*, and thus *f* (*r,N*) is maximum for 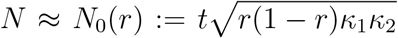. Then, one finds that 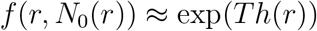, with 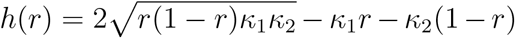, function which is maximum for 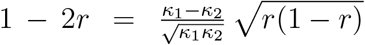. Then, noting that 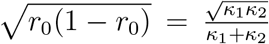 and 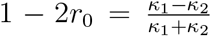. we obtain *r* = *r*_0_. After replacing *r* by *r*_0_, we find as expected *N*_0_ = *N*_0_(*r*_0_) ~ *κt*.

(2) Let us now carry out an expansion about that point in terms of x and y variables such that 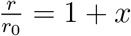 and 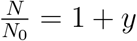 At first order in *x, y* when *x,y* → 0,

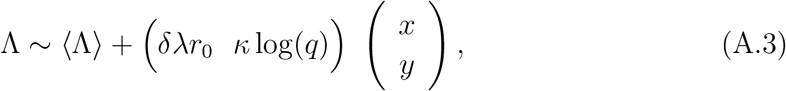

with *δλ* = λ_1_ – λ_2_. Let us write *f* (*r, N*) = *P* (*x,y*) with 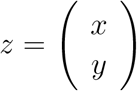. In the next point (3), we show that P is Gaussian with 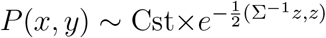 for a certain covariance matrix Σ. As a result:

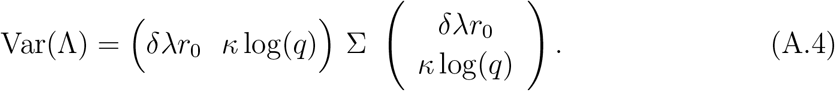

In the next point, we determine the matrix Σ.

(3) Let us perform an expansion to second order near *z* = 0,

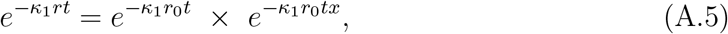

and similarly,

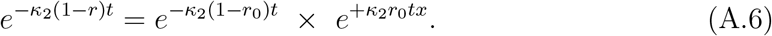

Then,

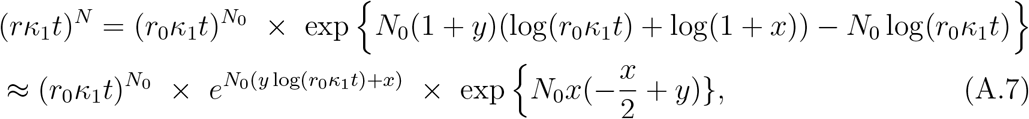

and similarly

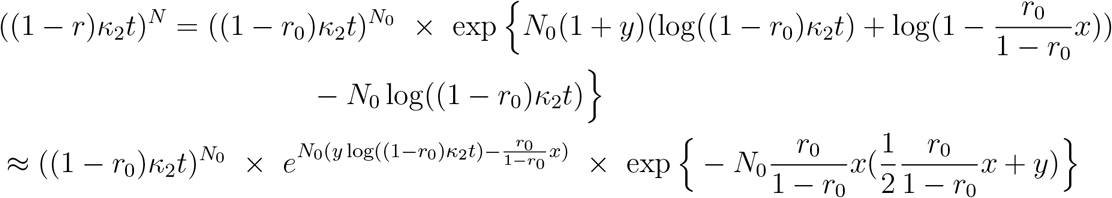

Then,

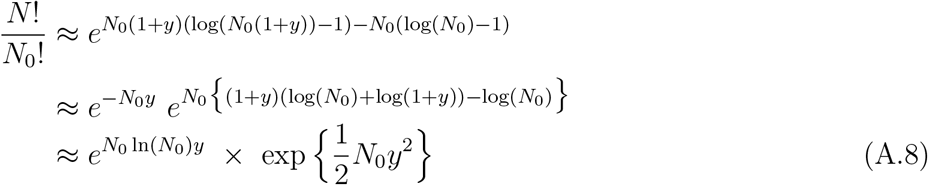

Taking the ratio 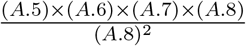, one checks immediately that the terms of first order in *x,y* cancel, and we get

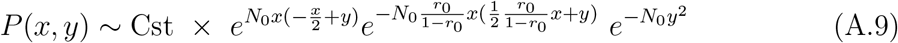

whence (using 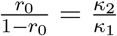)

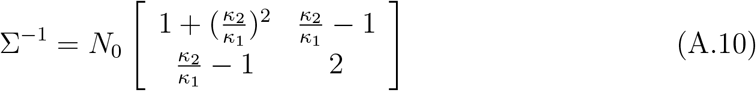

It is simple to show that det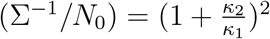, then

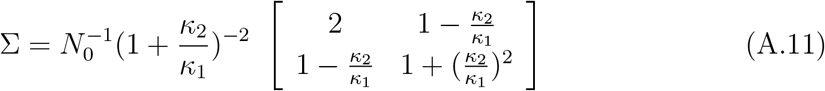

After evaluating (A.4) with this covariance matrix and replacing *r*_0_ by 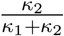, one finally obtains the result Eq. 21 for the variance in the KL approximation.

